# Free energy and kinetics of cAMP permeation through connexin26 hemichannel with and without voltage

**DOI:** 10.1101/2021.01.18.427208

**Authors:** Wenjuan Jiang, Yi-Chun Lin, Wesley Botello-Smith, Jorge Contreras, Andrew L. Harris, Luca Maragliano, Yun Lyna Luo

**Affiliations:** College of Pharmacy, Western University of Health Sciences, 309 E. Second St, Pomona, CA, USA; Department of Pharmacology, Physiology, and Neuroscience. New Jersey Medical School, Rutgers, The State University of New Jersey, Newark, NJ, USA; Department of Life and Environmental Sciences, Polytechnic University of Marche, Ancona, Italy; Center for Synaptic Neuroscience and Technology, Italian Institute of Technology, Genoa, Italy

## Abstract

The connexin family is a diverse group of highly regulated non-β-barrel wide-pore channels permeable to biological signaling molecules. Despite their critical roles in mediating selective molecular signaling in health and disease, the molecular basis of permeation through these pores remains unclear. Here, we report the thermodynamics and kinetics of binding and transport of a second messenger, adenosine-3’,5’-cyclophosphate (cAMP), through a connexin26 hemichannel. Inward and outward fluxes of cAMP were first obtained from 4 μs simulations with voltages and multiple cAMPs in solution. The results are compared with the intrinsic potential of mean force (PMF) and the mean first passage times (MFPTs) of a single cAMP in the absence of voltage, obtained from a total of 16.5 μs of multi-replica Voronoi-tessellated Markovian milestoning simulations. The computed transit times through the pore correspond well to existing experimental data. Both voltage simulations and milestoning simulations revealed two cAMP binding sites with binding constants and dissociation rates computed from PMF and MFPTs. The protein dipole inside the pore produces an asymmetric PMF, reflected in unequal cAMP MFPTs in each direction once within the pore. The free energy profiles under voltages derived from intrinsic PMF provided a unified understanding of directional transition rates with/without voltage, and revealed the unique role of channel polarity and the mobile electrolyte within a wide pore on the total free energy. In addition, we show how these factors influence the cAMP dipole vector during permeation, and how cAMP affects the local and non-local pore diameter in a position-dependent manner.

**Significance Statement:** Connexins are wide-pore channels permeable to cellular signaling molecules. They mediate molecular signaling crucial in physiology, pathology, and development; mutations in connexins cause human pathologies. However, the fundamental structural, thermodynamic, and kinetic determinants of molecular permeability properties are unknown. Using multiple molecular dynamics simulation techniques, we report, for the first time, an in-depth investigation of the free energy and the directional transition rates of an important biological signaling molecule, cAMP, through a connexin channel. We reveal the energetics and binding sites that determine the cAMP flux, and the effects of mobile ions and external electrical field on the process. The results provide a basis for understanding the unique features of molecular flux through connexins and other non-β-barrel wide-pore channels.

## Introduction

Connexin proteins form wide channels that mediate electrical and molecular signaling through cell membranes. They can function as plasma membrane channels (called “hemichannels) or as intercellular channels that allow direct transfer of small cytoplasmic molecules between cells. The intercellular channels (“gap junction channels”) are formed by end-to-end docking of two hemichannels across the extracellular gap between adjacent cells. The pores are relatively wide and are therefore permeable to atomic ions and small molecules in the size-range of key cellular signaling molecules including cAMP, cGMP, ATP, IP3, and glutathione. Each of the 21 human connexin isoform forms channels with distinct permeability and regulatory properties(*1, 2*). With the exception of electrical signaling in excitable tissues, the primary biological function of connexin channels is to mediate movement of small cytoplasmic signaling molecules between cells and/or to release them into the extracellular environment in a highly regulated manner. Mutations in connexins that alter channel function or expression produce human pathologies(*3, 4*). These mutations ultimately exert their pathological effects by disrupting the proper molecular permeability of junctional and plasma membranes that connexin channels mediate.

A large literature documents that channels formed by the different connexin isoforms have dramatically different permeability properties. Their unitary conductances range from 10 pS to 300 pS, their cation/anion permeability ratios (*P*_K+_/*P*_Cl-_) range from 8.0 to 0.8 and their permeabilities to fluorescent tracers are highly disparate. Strikingly, none of these parameters correlate with each other (e.g., the connexin channel with the largest unitary conductance is among the most size-restrictive). Permeabilities to biological signaling molecules are strikingly different among the different connexin channels, but are difficult to measure quantitatively, as the molecules are not fluorescent and do not carry significant current. Furthermore, for a given type of connexin channel, there are remarkable degrees of selectivity and relative permeability among biological permeants. These connexin-specific and permeant-specific permeability properties are not reasonably inferred from differences in permeabilities to fluorescent tracers(*5*). This suggests that there are, as yet unknown, structural/energetic determinants of molecular permeation that impart to each type of connexin channel specific, biologically required permeability properties. The underlying mechanisms for this are unknown, in spite of their clear biomedical and therapeutic importance. Investigation of these mechanisms by mutagenesis has not been informative in the absence of an understanding of the behavior, energetics, and interactions experienced by a molecule as it traverses the pore. Computational studies can provide the basis for this understanding.

We previously used Hamiltonian replica-exchange umbrella sampling to explore the energetics of uncharged permeant and non-permeant tracer molecules in the connexin 26 (Cx26) hemichannel(*6*). That study indicated that the determinants of molecular permeation differ from those that dominate the permeation of atomic ions through commonly studied ion-selective channels, emphasizing the unique aspects of small molecules with conformational and orientational degrees of freedom in a wide pore. The free energy and calculated relative permeabilities derived from that work were consistent with experimental findings. However, while validating the atomistic system and overall computational approach, the previous work did not provide quantitative kinetic information of the permeation process. Furthermore, the results of neutral tracer molecules do not characterize the biomedically crucial process of permeation by charged biological signaling molecules. The present study explores the binding and transport kinetics of a negatively charged second messenger, adenosine-3’,5’-cyclophosphate (cAMP), permeates a connexin pore.

A single permeation event with a timescale of sub-microsecond to microseconds is within reach of today’s computational power. However, a large number of transition events are required to obtain meaningful statistics, for which simulations of orders of magnitude longer than the mean transition time are required. One solution is to accelerate the permeation of charged molecules by imposing a voltage (cAMP carries a charge of −1e). Theoretically, if the system reaches a steady state under a constant electric field and maintains symmetric concentration on both sides of the channel, a mean flux rate can be estimated from the ensemble of nonequilibrium processes using a large number of permeation events. The accumulated density of cAMP along the channel axis during these events may provide an estimate of the locations of the energetic barrier(s) and binding site(s). Alternatively, one can choose an enhanced sampling method that is suitable for computing free energy and kinetics of the permeation process. In this study, we used both approaches to gain a comprehensive understanding of cAMP permeation with and without voltage, and in presence and absence of multiple permeants.

A molecular permeation rate is often dominated by the free energy profile or potential of mean force (PMF) along the channel lumen axis. Previously we estimated the relative transition rates between two sugar molecules using transition state theory (TST) based on PMF profiles(*6*). However, TST requires assumptions such as a single dominant transition state and no re-crossing at the barriers, which are often difficult to satisfy in complex biomolecular systems with rugged free-energy landscapes. To overcome this limitation, enhanced sampling techniques have been developed to calculate transition rates from molecular dynamics (MD) simulations. Of particular interest is the milestoning method introduced by Faradjian and Elber(*7*), which has been developed into several versions and used in many biophysical applications(*8*). Voronoi-tessellated Markovian milestoning is an implementation that allows reconstruction of the long-time dynamics of a system from independent simulations confined within a set of cells spanning the space of the reaction coordinates(*9*). This method has successfully captured the rates of CO entry/exit in myoglobin(*10*), and recently the ligand binding kinetics(*11*). Here, we make use of the “soft-walls” version, which confines the sampling within the Voronoi cells using flat-bottom harmonic restraining potentials(*12*). This approach is easy to implement, allowing us to take advantage of CUDA accelerated MD packages, and has been shown to yield the same results as the original “hard-walls” version, which instead inverts atomic velocity at the cell boundaries. The “soft-walls” version has been used to study nucleation of an ionic liquid(*13*) and ion permeation across a claudin-15 paracellular channel(*14*). Here we adopt this approach to explore the permeation of cAMP through a Cx26 hemichannel.

## Results

### cAMP permeation under opposite voltages

We first investigated inward/outward cAMP permeation rates under voltages in the presence of symmetric cAMP concentrations. To obtain a sufficient number of transition events and ensure unidirectional flux, we applied +/-200 mV transmembrane potentials to accelerate permeation. 27 cAMP molecules and 27 Mg^2+^ ions, corresponding to 26.5 mM, were present on each side of the membrane. To mimic physiological salt concentration and also to neutralize the protein charge, 82 K^+^ and 163 Cl^-^ ions were added to the bulk aqueous compartments, corresponding to 80.5 mM K^+^ and 160.1 mM Cl^-^ (**Table S1**). Using z<∣50∣ Å as boundaries of the hemichannel (center-of-mass of protein is at z=0 Å), 9 single-molecule transitions were observed at +200 mV and 12 transitions at −200 mV during each 2 μs simulation (**Figure 1, Table 1**, and raw data in **Table S2**). At each voltage, all transitions were in the same direction.

**Table 1.** Transition time of cAMP through C×26 hemichannel under voltage.

|  | +200mV | -200mV |
| --- | --- | --- |
| cAMP flux direction | Inward | Outward |
| Transition events in 2 μs | 9 | 12 |
| Events per μs | 4.5 | 6.0 |
| Time between transition events | 222 ns | 166 ns |
| Mean transition time (CI<sub>95</sub>) * | 448 ns (204-1646) | 510 ns (249-1571) |
| Mean dwell time (CI<sub>95</sub>) | 305 ns (149-940) | 398 ns (205-1086) |
| Mean barrier crossing time (CI<sub>95</sub>) | 143 ns (79-330) | 106 ns (54-290) |

\*Sample mean and 95% confidence intervals are based on Maximum Likelihood Estimate by fitting the exponential distributions. See code at https://github.com/LynaLuo-Lab/MD-data-uncertainty-analysis/blob/master/confidence_interval_exponential.ipynb.

**Figure 1.**
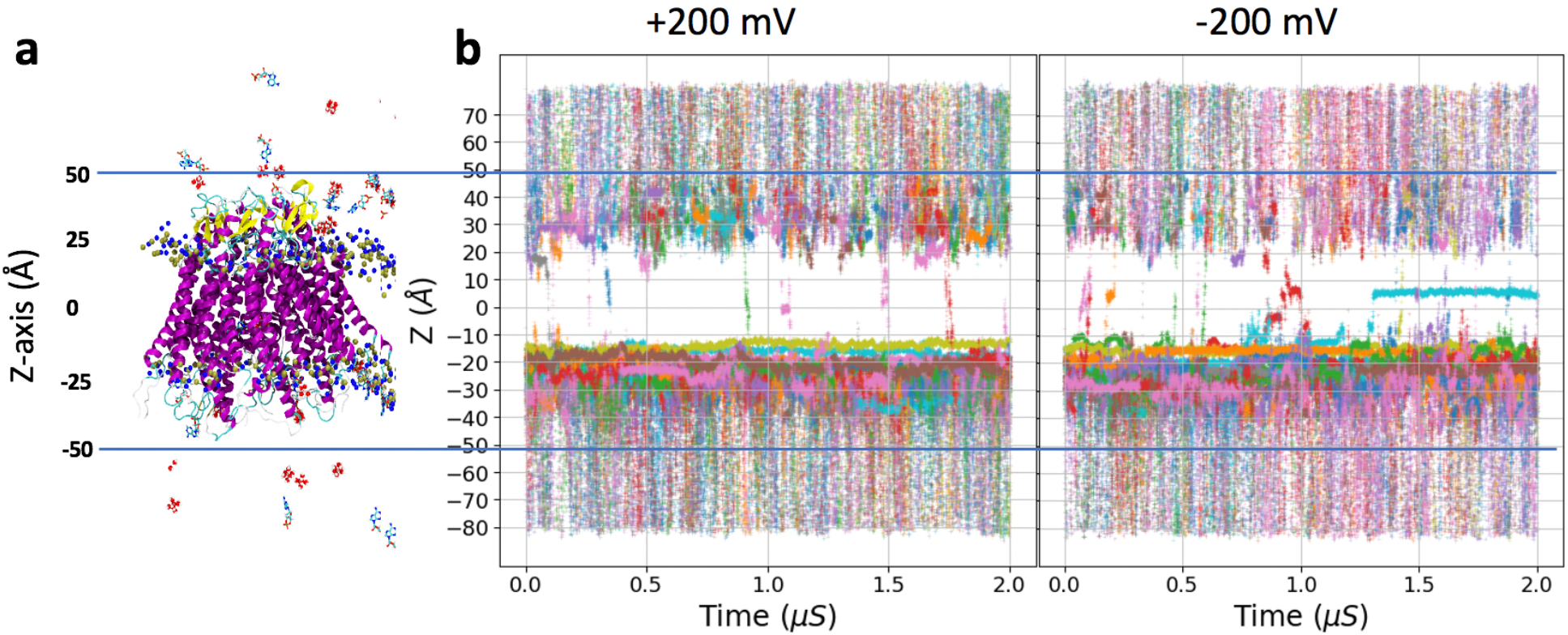
Simulations of cAMP permeation through C×26 at +200 mV and −200 mV membrane potential. **a**. Snapshot of the simulated system. Protein backbone is shown in cartoon mode and colored by the secondary structure (helix in magenta, beta-sheet in yellow, disordered loop in cyan). cAMP molecules are shown in licorice with atoms colored (red oxygen, cyan carbon, blue nitrogen, yellow phosphate). Lipids, ions, and water molecules are not shown. **b**. The z-coordinates of all 27 cAMP molecules in the system during the simulations are shown in different colors. Note that the flux of Camp is in the direction opposite to the field; thus, cAMP flux is inward (down in this figure) under +200 mV and outward under −200 mV.

To estimate the uncertainty due to the small number of events, we calculated the confidence interval for the average transition time (***<t>***) of cAMP through the channel by fitting the transition times to an exponential distribution. Based on a maximum likelihood estimation, MFPT is 448 ns at +200 mV with 95% confidence interval (CI_95_) of 204-1646 ns, and 510 ns with CI_95_ of 249-1571 ns at −200 mV. The average time between consecutive transition events (**τ** = length of simulation/number of transits) is 222 ns at +200 mV and 166 ns at −200 mV. Given the confidence intervals, there is no significant difference in the flux in each direction. The ratio ***<t>***/**τ** is 2.0 and 3.1, for +200 mV and −200 mV, respectively, which indicates there are on average two to three cAMP molecules in the channel at any given time at each voltage. Inspection of the trajectories shows that this is only due to the accumulation of cAMP molecules at the intracellular entrance (**Figures 2 and 3**).

**Figure 2.**
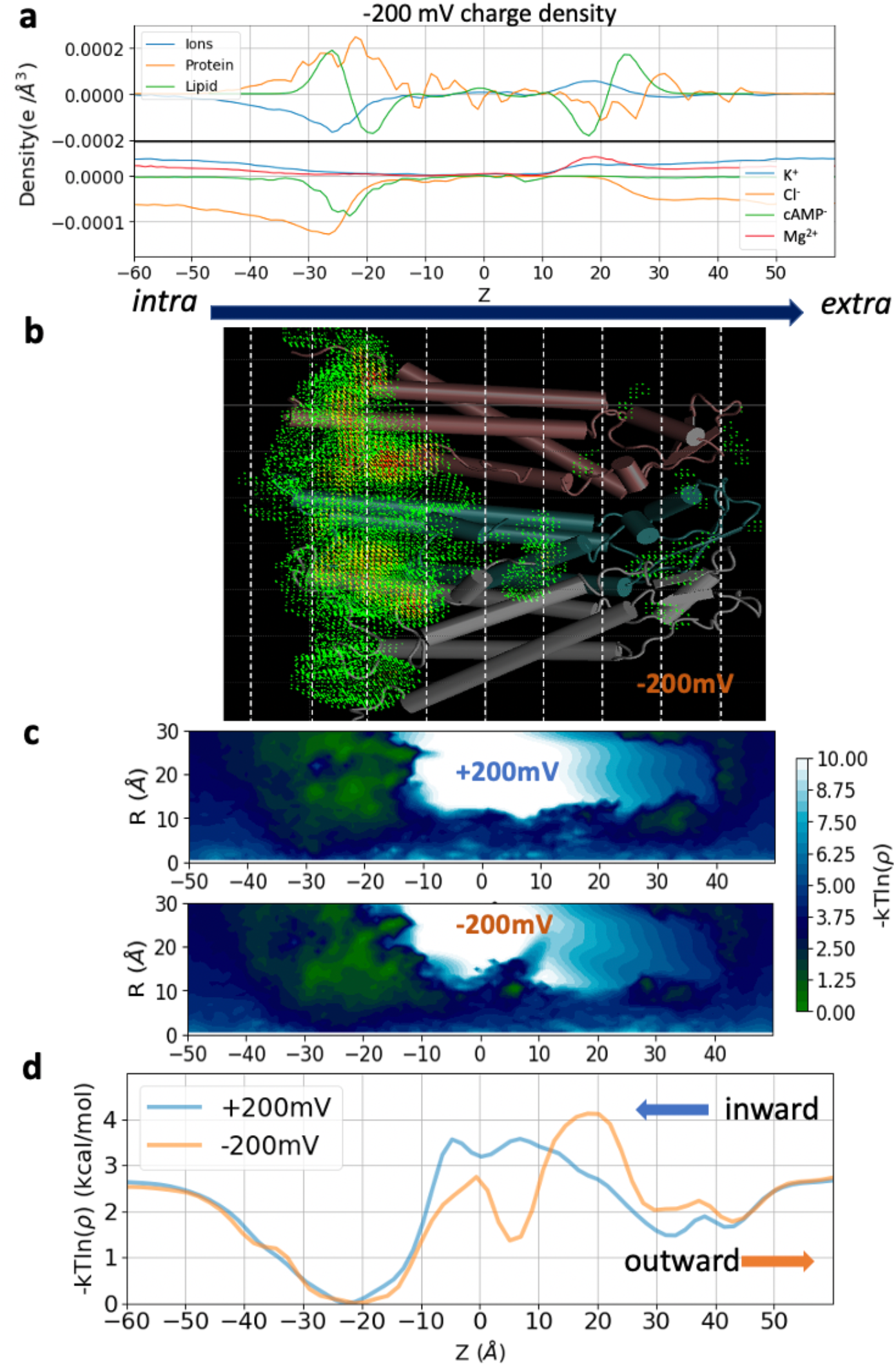
Charge density profile and cAMP log-density profiles in 3D, 2D, and 1D. **a**. Charge density profiles of protein, lipids, total ions and individual charges groups (K^+^, Cl^-^, cAMP^-^, Mg^2+^) along the channel z-axis obtained from the −200 mV trajectory. **b**. Volumetric map of cAMP 3D density under −200 mV simulation. **c**. Boltzmann inversion *-kTln(p(z))* of the accumulated cAMP density profile in 2D along the channel z-axis and radial axis R. **d**. 1D log-density plot of the cAMP density profile within the pore along the channel z-axis. Arrows indicate the direction of cAMP flux at each voltage. Note: **a** represents charge density in the entire simulation system, while **b, c**, and **d** depicts the cAMP density only within the pore using the same cylindrical radius cutoff of 30 Å as in the milestoning simulation below.

**Figure 3.**
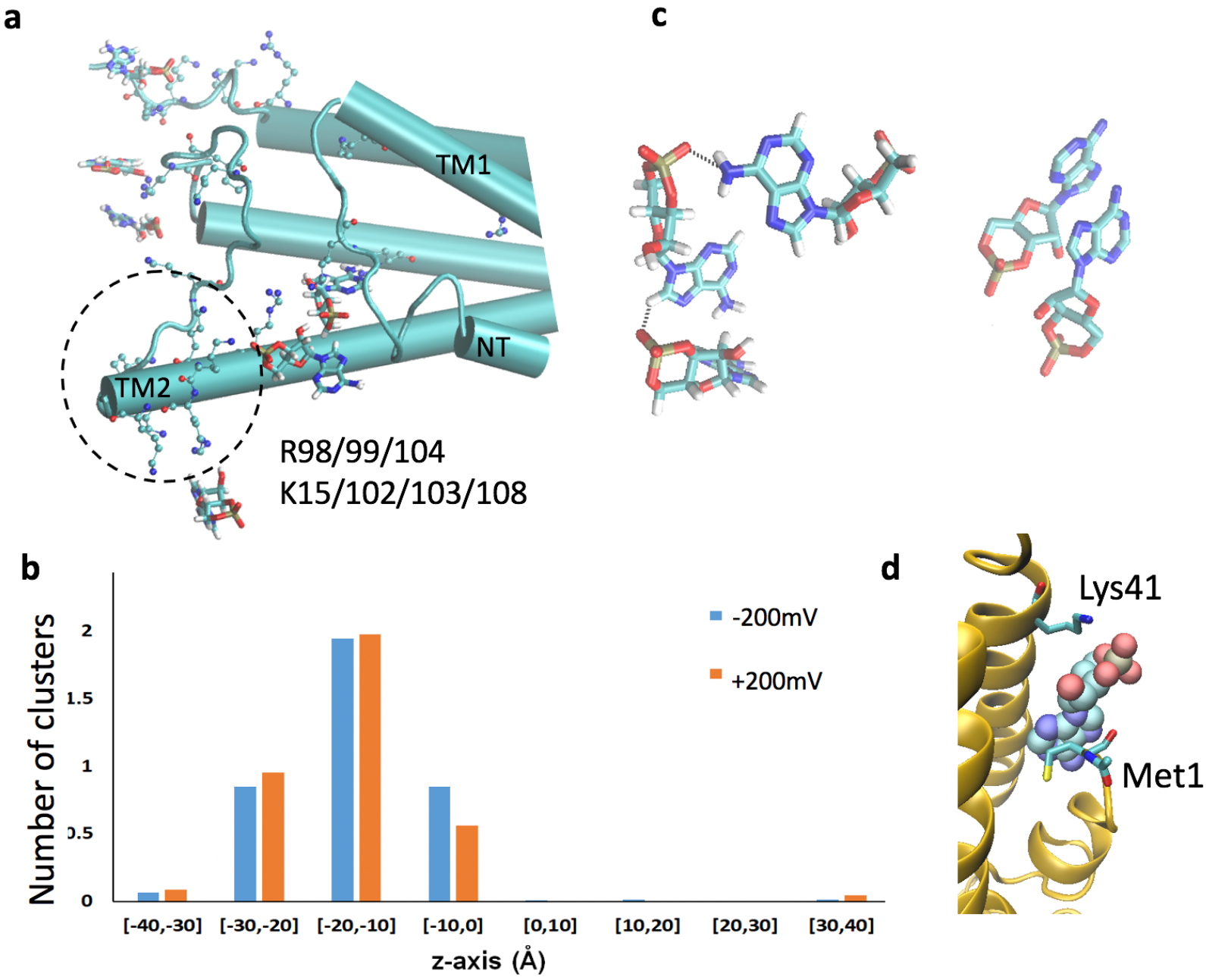
Permeant-permeant interactions and permeant-channel interactions. **a**. Snapshot showing cAMP molecules (in licorice) near positively charged residues (in CPK mode) at the intracellular entrance. Protein is in cartoon mode. **b**. Number of cAMP clusters in each 10 Å window during +/-200 mV simulations. A cluster is defined as at least two cAMP molecules with centers of mass within 20 Å at any time during the simulations. **c**. Snapshots of two cAMPs clustered via pi-stacking and three cAMPs clustered via hydrogen bonds. **d**. Snapshot showing one cAMP trapped between Lys41 and Met1.

Although the experimental transition time of cAMP under voltage is not available, a cAMP/K^+^ permeability ratio of 0.027 was reported using simultaneous measurements of Cx26 junctional conductance and reporter-based intercellular transfer of cAMP(*15*). We calculated K^+^ transition time in presence of 27 cAMP is 13.1 ± 13.0 ns at +200 mV and 6.8 ± 7.8 ns at −200 mV. If we assume the time needed to cross the junctional channel (two hemichannels docked at the extracellular ends) is the sum of the transition time in two opposite directions, we obtain a cAMP/K^+^ ratio of 0.021, reasonably close to the experimental ratio.

### Pore cAMP density profiles from voltage simulations

The density profiles accumulated from voltage simulations can provide an estimate of the locations of energetic barriers and binding sites. The charge densities of protein, lipids, total ions, and individual ions (K^+^, Cl^-^, cAMP^-^, Mg^2+^) are plotted along the channel z-axis (**Figure 2a**). It can be seen that the channel is largely positive at intracellular entrance where the highest density of Cl^-^, cAMP^-^ ions are located. The accumulation of cAMP at the intracellular entrance can also be seen clearly in **Figure 1**. A volumetric map showing the 3D density of cAMP inside the channel from the −200 mV simulation is illustrated in **Figure 2b**. A smooth indicator of cAMP distribution is shown using Boltzmann inversion (−*kTl n*(ρ)) of the cAMP density; we will call these *log-density plots*. **Figure 2c** shows the 2D cAMP log-density along the z-axis and the radial axis *R*. **Figure 2d** shows the 1D cAMP log-density within the pore along z. Note that these density profiles are acquired from nonequilibrium simulations under the influence of voltage. They indicate how the cAMP behaves under the influence of both external voltage and the intrinsic free energy, hence do not represent the equilibrium free energy profiles. The symmetric cAMP density in the bulk regions on each side of the channel is the result of the periodic boundary condition used for MD simulations.

The 1D log-density profiles (**Figure 2d**) have a broad minimum at the intracellular entrance of the channel and a major peak in the middle of the channel. The shapes of the global minimum region are essentially identical under both voltages. There is a second well in the −200 mV density that appears to bisect the central peak. The dwell times of cAMP (**Table 1**) at the broad minimum (−50<z<10 Å) are quite large under both voltages: 305 ns at +200 mV, and 398 ns at −200 mV (**Table 1**). In contrast, the rest of the channel region (−10<z<50 Å), which contains the major barriers, was crossed by cAMP more rapidly: 143 ns for inward flux (at +200 mV), and 106 ns for outward flux (at −200 mV).

The broad binding well (−50<z<-10) present at both voltages is located in the C-terminal regions of the second transmembrane helix (TM2) and the N-terminal helix (NTH). Contact frequency analysis between cAMPs and protein sidechains indicates that cAMP bind to R99/104 and K103 on TM2 over 70% of the simulated time, and bind to K102 and R98 on TM2, and K15 on NTH over 30% of the time (**Figure 3a**), thus providing a basis for the cAMP accumulation at this region. Further investigation revealed that the cAMP forms clusters between −30<z<0 Å, while almost no clusters are found in the rest of the channel (**Figure 3b**). Most of the clusters contain 2 cAMPs, which interact through pi-stacking of adenosine rings. Clusters of 3 cAMPs forming hydrogen bonds with each other also exist (**Figure 3c**). These clusters are not seen in the bulk. It appears that the positively charged residues at the intracellular entrance of the pore facilitate cAMP clustering by reducing the translational and rotational entropy of the molecules. To check whether this accumulation produce a “crowding” effect on entry into the pore, we compared the unoccupied lumen radius with and without cAMP molecules present. **Figure S1** suggests that the accumulation of cAMP had little effect on the average available cross-sectional area at the pore entrance. In addition, the peaks in the two log-density profiles (**Figure 2d**) do not closely correlate with the narrow regions of the pore (see **Figure S1 and Figure 8a** for two different radius measurements), indicating that steric hindrance is not a major contributor to the cAMP transition barrier.

During one of the 12 permeation events at −200 mV simulation, one cAMP was trapped between Met1 in the N-terminal helix (NTH) and Lys41 on the first transmembrane helix (TM1) (**Figure 1b, 3d**). This was reflected in the density profile as a large dip in the peak at z=5 Å (**Figure 2d**). Both NTH and Lys41 have been suggested to be involved in voltage-sensing in Cx26(*16, 17*). This raised the question of whether the instance of a long residency of a cAMP molecule at this particular position at −200 mV, but not evident at +200 mV or 0 mV (from PMF milestoning below) is a consequence of voltage-driven repositioning of these charged moieties. To evaluate their responses to the local electric field, we plotted the angles between the z-axis and the principal vector of Lys41 or NTH (residues 1 to 11) during 2 μs simulations at +/-200 mV (**Figures S2a and S2b**). Except for a clear reduction in Lys41 fluctuation in subunit 5, where the cAMP was trapped, there is no clear preference in the orientation of the Lys41 or NTH in response to the two opposite voltages. Therefore, this trapped cAMP is unlikely to be due to the effect of voltage on the protein. Of course, this result does not indicate that the NTH and Lys41 are uninvolved in voltage-sensing, only that they did not respond to +/-200 mV within the 2 μs simulations.

### cAMP permeation free energy using Markovian milestoning

The simulations above provide a nonequilibrium view of the cAMP transition process driven by voltage. Two opposite voltages resulted in similar cAMP density profiles and similar ranges of flux rates at each voltage. However, the small number of stochastic events (21 in total) led to the large uncertainty in the mean passage time of the transition event. In contrast to a long trajectory exploring the whole channel, multiple MD simulations confined in intervals partitioning the space can offer sufficient statistics within a shorter running time. Here, we used Voronoi-tessellated Markovian milestoning MD simulations (hereafter referred to as milestoning simulation) on a tessellation along the z-coordinate of the center-of-mass of a single cAMP to estimate the PMF and the kinetics of permeation through the channel at zero membrane voltage.

**Figure 4** shows the 1D PMF (in red) obtained from the equilibrium probability of finding cAMP in each milestoning cell. This PMF of a single cAMP permeating without applied voltage has features similar to the log-density plots of the accumulated density of cAMP from the voltage simulations. A free energy barrier spans between −20<z<30 Å, with the peak located at z = +20 Å, the same as the peak in log-density from the −200 mV simulation. The height of the barrier relative to bulk (1.2 kcal/mol) is slightly lower than in −200 mV log-density (1.8 kcal/mol) and higher than in +200 mV (0.8 kcal/mol). The major well on around z = −20 Å from voltage simulations are much broader and more favorable (−2.8 kcal/mol relative to bulk) than the well in the PMF (−2 kcal/mol), likely due to the cAMP clustering, which is absent in the single cAMP milestoning. The dip that split the peak at z = +5 Å under −200 mV also shows up in the PMF, but with much smaller magnitude. It is thus possible that the magnitude of the dip at −200 mV in the log-density plot is overestimated due to the contribution of the single, rare event of cAMP trapping (evident in **Figure 1b**).

**Figure 4.**
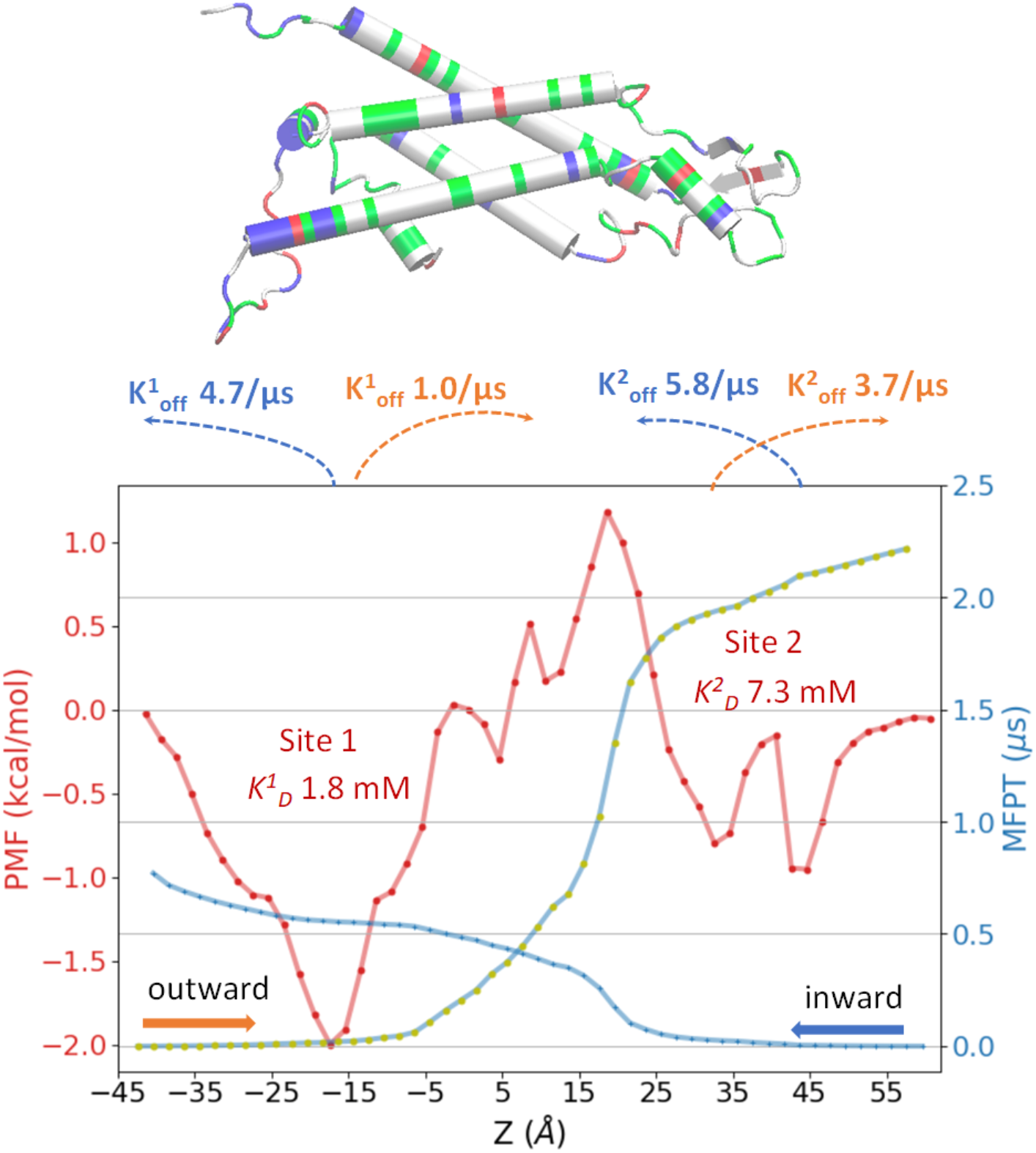
Free energy and kinetics of a single cAMP permeation through Cx26 hemichannel from milestoning simulation (V=0 mV). Potential of mean force (PMF) in red, inward mean first passage times (MFPT) in blue, and outward MFPT in yellow dotted blue line. Binding constants and dissociation rates derived from PMF and MFPT profiles are indicated. The graphic at the top shows the backbone of one Cx26 subunit with z-positions aligned with the plot below (basic residues in blue, acidic in red, polar in green, and nonpolar in white).

Similar to the voltage simulations, milestoning simulation without voltage reveals two binding sites for cAMP, a major one (site 1) near the intracellular entrance between NT and TM2 (−2 kcal/mol relative to bulk value) and a (bisected) much smaller one (site 2) at the extracellular loop (E1) region (−0.7 kcal/mol relative to bulk). The single cAMP dissociation constant, *K*_*D*_, can be estimated from the single cAMP equilibrium PMF using Eq. 1:

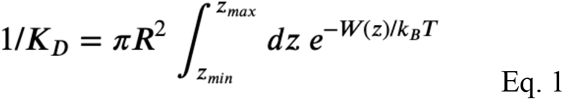

*W*(z) is the PMF with bulk as reference. *R* is the radius of a cylindrical restraint (30 Å). Integrals over individual energy wells indicate that cAMP will bind to the intracellular site (binding site 1: −43.4<z<18.6 Å) with *K*_*D*_ of 1.8 mM and to the extracellular site (binding site 2: 18.6<z<60.6 Å) with *K*_*D*_ of 7.3 mM (**Figure 4a**). The integral over the entire PMF yields a total *K*_*D*_ of 1.4 mM for the channel. Thus, at the bulk cAMP concentration of the voltage simulations (26.5 mM), both binding sites are likely highly occupied, while at the concentration equivalent to the single cAMP used in milestoning (1 mM), none of the binding sites would be occupied more than 50%.

### cAMP permeation kinetics using Markovian milestoning

According to the MFPT profiles in **Figure 4**, from the first to the last milestone at the boundaries of the channel, it takes about 0.77 μs for inward flux and 2.22 μs for outward flux. Thus, the inward flux is 2.9 times faster than outward flux once a single cAMP enters the channel. The faster inward than outward flux is the direct consequence of the asymmetric channel, reflected in the asymmetric PMF showing a maximum of 3.2 kcal/mol barrier for outward flux, which takes 1.0 μs to cross, while the two smaller barriers of 2.0 kcal/mol for inward flux only take 212 ns (site 1) and 145 ns (site 2) to cross. MFPT profiles can provide direct information about the dissociation rate (k_off_) of a single cAMP. For outward permeation, k_off^1^_ for binding site 1 is 1.0 μs^-1^ (−17.4 <z<18.6) and k_off^2^_ for binding site 2 is 3.7 μs^-1^ (32.6 <z<60.6). For inward permeation, k_off^1^_ is 4.7 μs^-1^ (−41.4 <z<-17.4) and k_off^2^_ is 3.7 μs^-1^ (18.6 <z<42.6). With the equilibrium constant *K*_*D*_ from PMF, we can also estimate the association rate k_on^1^_of 0.6 μs^-1^mM^-1^ and k_on^2^_of 0.5 μs^-1^mM^-1^ for outward permeation, and k_on^1^_of 2.6 μs^-1^ mM^-1^ and k_on^2^_of 0.8 μs^-1^ mM^-1^ for inward permeation.

It should be noted that the MFPT is subjected to the condition that the channel is occupied by only one permeating molecule at any time. Thus, it only represents the transit time of a single permeant traversing an otherwise permeant-free channel. The full kinetics and flux at finite bulk concentrations of the permeant also depend on bulk diffusivity and concentration, and the diameters of the entrance at each end of the pore. For instance, diffusion current to a disk-like adsorber is ***I=4DRC***, where ***C*** is the permeant concentration in the infinite bulk, ***D*** is bulk diffusion constant, and ***R*** is the radius of the disk-shaped absorber(*18*). Interestingly, for Cx26 hemichannel, the intracellular entrance (radius ***R***=25 Å) is larger than the extracellular entrance (***R***=10 Å) (see pore radius in Figure 8a). Taking into account the effective radius of cAMP as *r*=3 Å (calculated from radius of gyration), the effective pore radii are 22 Å and 7 Å for the intracellular and extracellular pore entrances, respectively, indicating that it is ∼3 fold more likely for cAMP to reach the pore by random diffusion from bulk to the intracellular than the extracellular side.

Quantitative experimental measurements of the flux of cAMP through Cx26 channels in cells are very complex and subject to a variety of potential confounding factors (*5*). Two studies, which used different indirect strategies to report cAMP flux through junctional channels in the absence of junctional voltage, yielded estimates of cAMP permeability that differed by nearly a factor of 8 (6.2 and 47 × 10^−3^ um^3^ sec^-1^) (*15, 19*). Using the volume of the Cx26 channel obtained from the simulations, these permeability predict cAMP transit rates through junctional channels of 14 μs and 1.9 μs, respectively, which bracket the ∼3 μs transit time inferred from our studies (sum of hemichannel “outward” and “inward” transit times of 0.77 and 2.22 μs).

### Influence of voltages on free energy profile

The milestoning simulations at zero voltage yielded an inward transition time is 2.9 times faster than the outward transition time (0.77 vs 2.22 μs). Interestingly, the mean inward/outward barrier crossing time under voltages are quite similar: 143 ns (CI_95_ 79-330 ns) at +200 mV, and 106 ns (CI_95_ 54-290 ns) at −200 mV (**Table 1**). Below we show that this is likely due to the negative voltage reducing the free energy barrier for cAMP outward flux.

It has been shown previously that the total PMF under voltage *W*_*tot*_(*z*) can be computed as the sum of the intrinsic PMF in the absence of external field *W*_*eq*_(*z*) (in this case the equilibrium PMF derived from milestoning), and the additional potential introduced by the external field *qδϕ*(*z*) (Eq. 2)(*20-22*). This additional potential has two components. One is the constant electric field throughout the entire simulated periodic cell *E= V*(*z*)*/L*_*Z*_, where *V*(*z*) is the voltage linear to *L*_*Z*_, the length of the PBC box in the *z*-direction. The other component is the reaction potential due to the voltage-induced changes of the spatial distribution and orientation of the water dipole and mobile ions, as well as flexible and charged atoms in the protein and membrane(*23*). This approach presumes that the channel and the permeant do not undergo substantial conformational changes due to the external field within the time of simulation (2 μs in this case). This assumption is supported by our results from +/-200mV simulations showing highly similar cAMP density distribution (**Figure 2**) and pore radius profile (**Figure S1**).

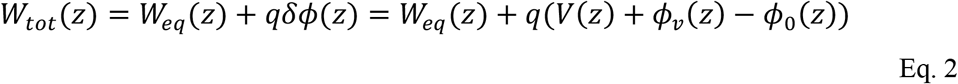

The reaction potential introduced by external field may be approximated by the difference in electrostatic potential in presence and absence of the external potential, *ϕ*_*v*_(*z*) − *ϕ*_0_(*z*). To calculate this difference in electrostatic potential, three additional 100 ns simulations at −200, 0, +200 mV were carried out for the same system but without cAMP. 3D electrostatic potential maps *ϕ*(r) were calculated based on all charged atoms in the simulated system ρ_*i*_(*r*) by solving the Poisson equation 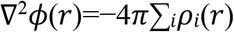 (*r*) on a 1 Å resolution grid using the VMD PMEPot plugin(*24*). **Figure 5a** and **5b** show the 1D potential along the central pore z-axis without voltage (in blue) and under +/-200 mV (in orange).

**Figure 5.**
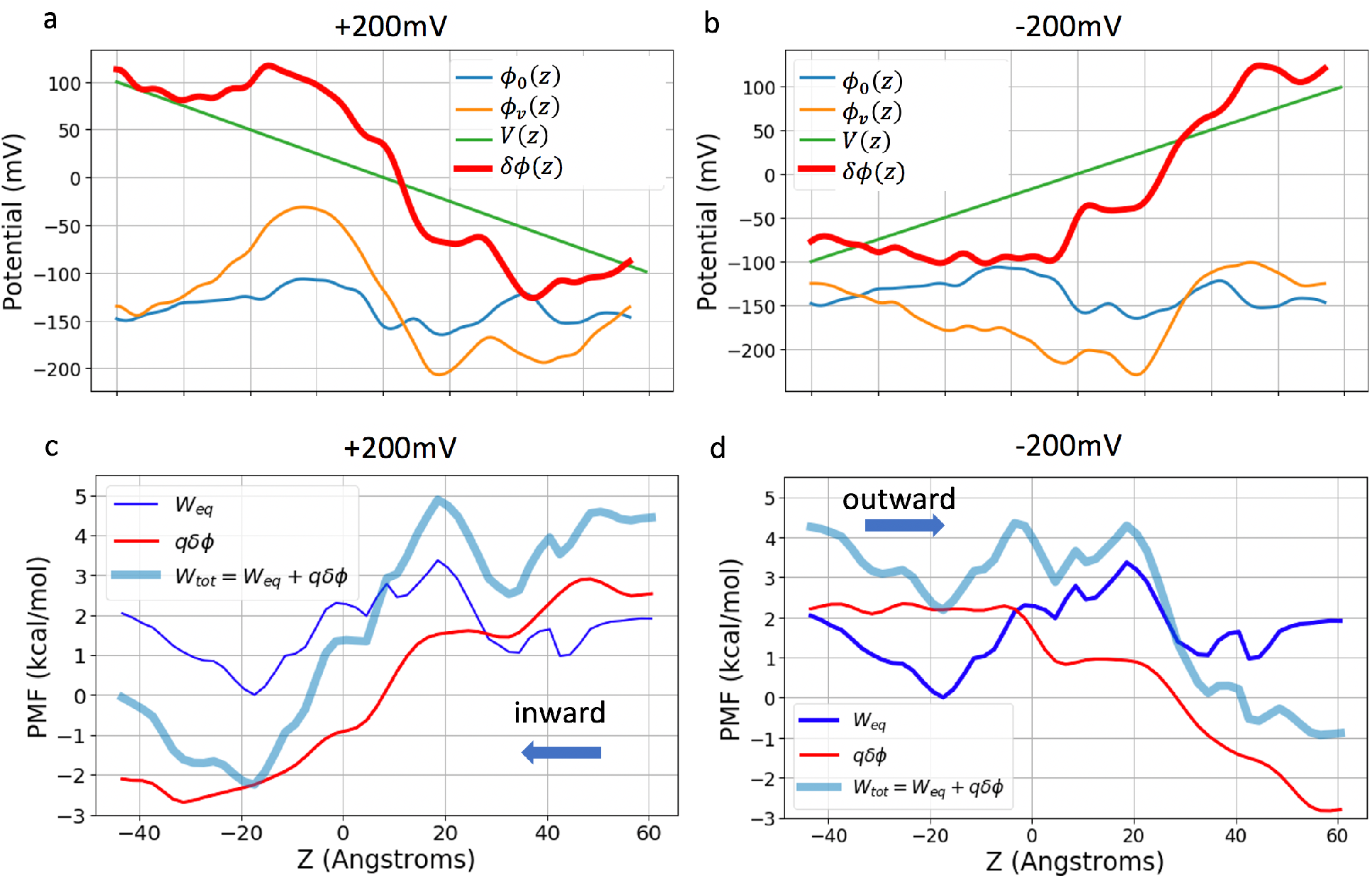
Electrostatic potentials and PMFs under voltages. **a**,**b**: applied potential in green (+/-200 mV) and the electrostatic potential under voltage *ϕ_v_*(z) in orange, and without voltage *ϕ_0_*(z) in blue. The sum of applied potential and reaction potential (see Eq. 2) in shown in red. **c**,**d**: Total PMF under a constant electric field, W_tot_(z), from the intrinsic PMF in absence of an electric field W_eq_(z), and the additional electrostatic potential energy introduced by the applied field q**δ***ϕ*(z).

Cx26 is largely positive in the intracellular side (−40<z<0), negative near the extracellular side (15<z<25), and slightly positive near extracellular entrance (25<z<35) (**Figure 2a**), thus it has an overall dipole vector pointing towards the positive z-direction. Consequently, within the protein-membrane region, positive voltage produces a potential *ϕ*_*v*_(*z*) that enhances the protein dipole, while the one from negative voltage counters the protein dipole (**Figure 5ab** orange lines). In the bulk region and lipid headgroup region (z > ∣15∣), the *ϕ*_*v*_(*z*) nearly cancels out the external field *V*(*z*). The sharp drop of the *ϕ*_*v*_(*z*) under +200 mV can be visualized on 2D-electrostatic potential maps of the whole system (**Figure 6a**), and the 3D-electrostatic potential overlaid onto the solvent mass density iso-surface **(Figure 6b**) or onto the cAMP density iso-surface (**Figure 6c**).

**Figure 6.**
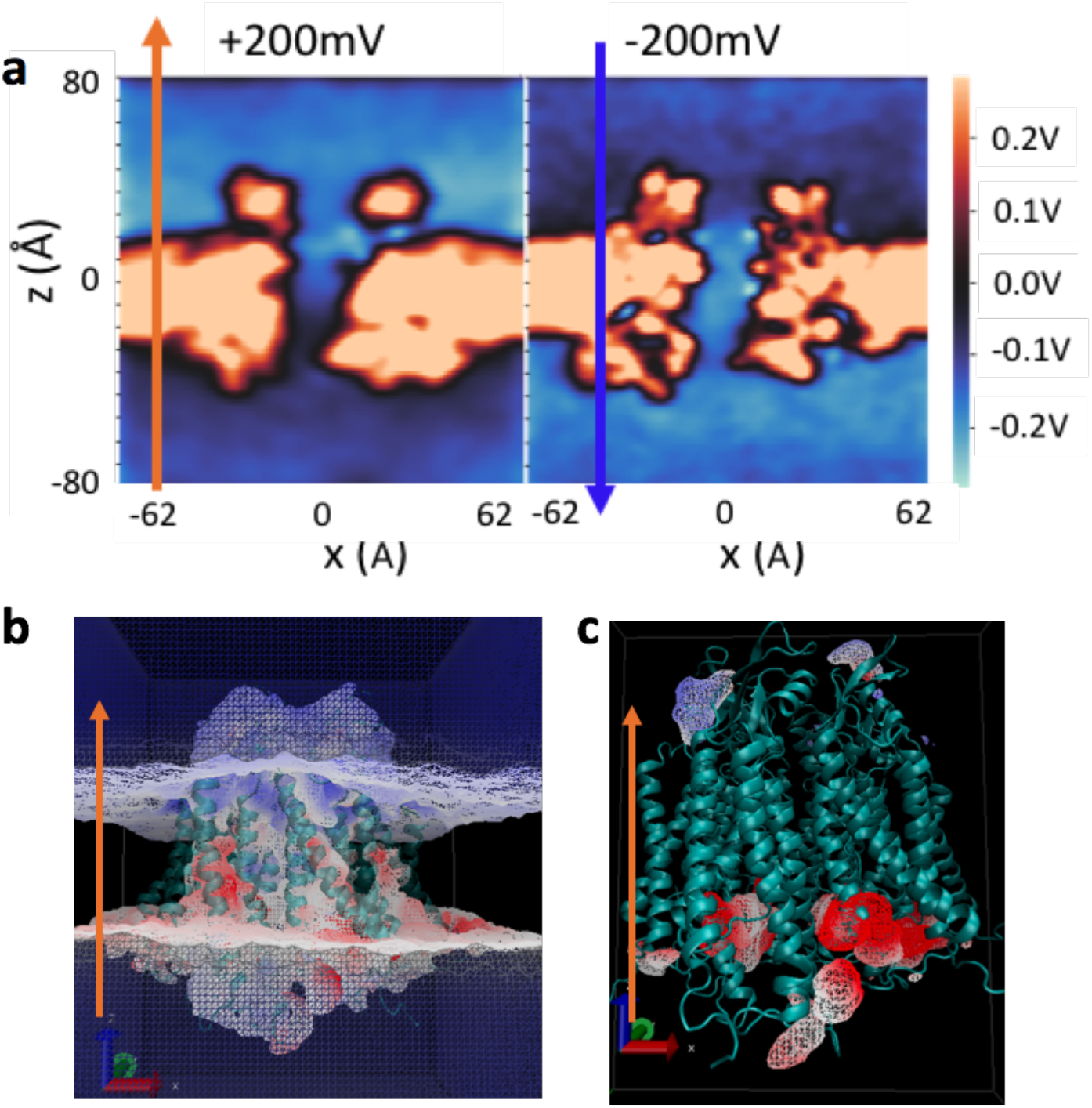
Electrostatic potential at +200 mV and −200 mV membrane potential. **a**. 2D-electrostatic potential maps (vector 1,0,1 in Cartesian space) at +/-200 mV voltage. **b**. The 3D-electrostatic potential at +200 mV overlaid onto the solvent mass density iso-surface. The color scale is −12.11 to 32.95 *kT*/e (−323 to +880 mV) from blue to red. Only the electrostatic potential at +200 mV is shown here as the color change in −200 mV is much less prominent (due to the opposite orientation of the intrinsic dipole as described in the text). **c**. Electrostatic potential overlaid onto the cAMP density iso-surface at +200 mV. The color scale is −267 to +267 mV. The iso-surface contour cutoff is 0.1 amu/Å^3^ for panels b and c. All data in this figure are calculated from the 3D electrostatic potential map *ϕ*(*r*) based on all charged atoms in the simulated system by solving Poisson’s equation on a 1 Å resolution grid using the VMD PMEPot plugin. PMEPot approximate point charge by a spherical Gaussian with an Ewald factor of 0.25. *ϕ*(*r*) is reported as the average of 1000 snapshots from the last 200 ns.

**Figure 5 cd** show the total PMF under each voltage *W*_*tot*_(*z*) obtained from Eq. 2. Clearly, +200 mV facilitates the inward flux of the negatively charged cAMP by increasing the free energy on the extracellular side of the protein (z>0) and decreasing the free energy on the intracellular side (z<0). The two inward flux energy barriers remain similar to those of the intrinsic PMF (∼2.3 kcal/mol). However, the outward *W*_*tot*_(*z*) at −200 mV significantly reduced the outward barrier from 3.2 kcal/mol of the intrinsic PMF to 2.3 kcal/mol. Thus, the asymmetry of the inward *vs*. outward MFPTs of cAMP within the pore at zero voltage is eliminated by the external voltage. This “voltage-equalizing” effect of permeation kinetics is a unique feature of large-pore channels that have a protein dipole and mobile electrolytes inside the pore.

### cAMP dipole orientations during transit with and without voltage

cAMP is a fairly rigid molecule with a dipole moment of 37.1 Debye. **Figure 7a** shows the probability distribution of the angle between the cAMP dipole vector and the z-axis from milestoning simulations. Strikingly, the cAMP molecule rotates nearly 180 degrees five times on its way through the pore (−40∼-20, −20∼0, 0∼10, 10∼20, and 20∼40 Å along the z-axis, also see **Video1**). To understand this, the z-components of the force vector acting on the cAMP from the rest of the system (protein, water, ions, lipids) are decomposed into electrostatics and vdW terms and plotted for each milestoning cell (**Figure 7b**). The reciprocal forces of electrostatics and vdW terms along the channel indicate that cAMP has close interaction with the walls of the pore lumen. The positive forces push cAMP in the positive z-direction, thus facilitating outward permeation, while the negative force facilitates inward permeation towards the negative z-direction. The cAMP dipole rotates as the electrostatic force vector switches the sign. Thus, these re-orientations of cAMP dipole vector in absence of external voltage are due to the local electric field along the pore lumen.

**Figure 7.**
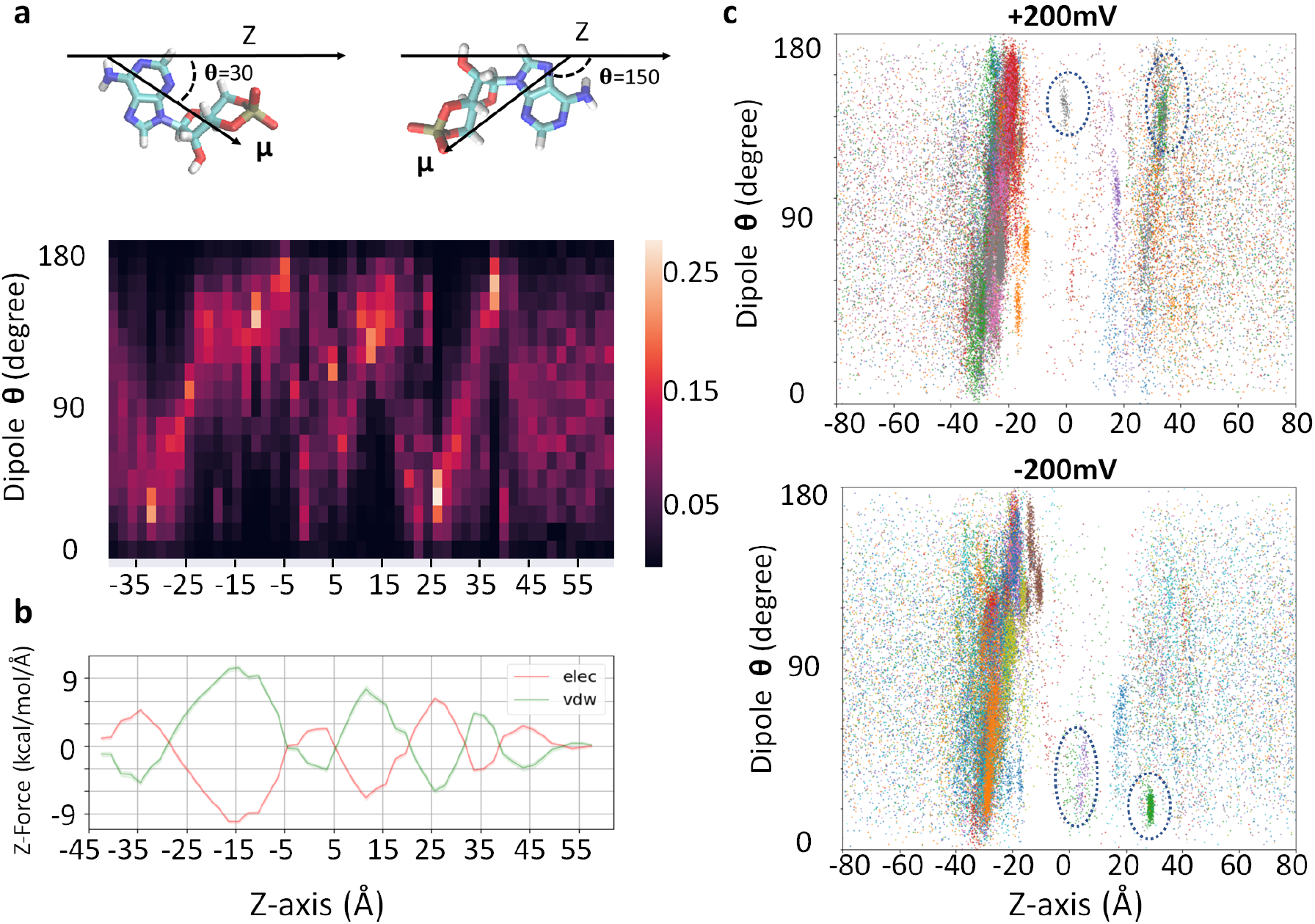
Dipole moment distribution and force decomposition for cAMP molecules along z-axis (Å). **a**. cAMP dipole angles are shown as a probability distribution for each milestoning simulation cell along the z-axis. Color bar shows the probability scale, lighter color represents higher probability. Representative dipole angles are illustrated on the cAMP molecule above. **b**. Mean electrostatic and vdW forces along z-axis on cAMP. Lines represent the running average of three milestoning cells and shaded areas represent standard error of mean within each milestoning cell. **c**. Dipole angle scatterplot of permeating cAMP at − 200 mV and +200 mV voltage simulations, respectively.

**Figure 8.**
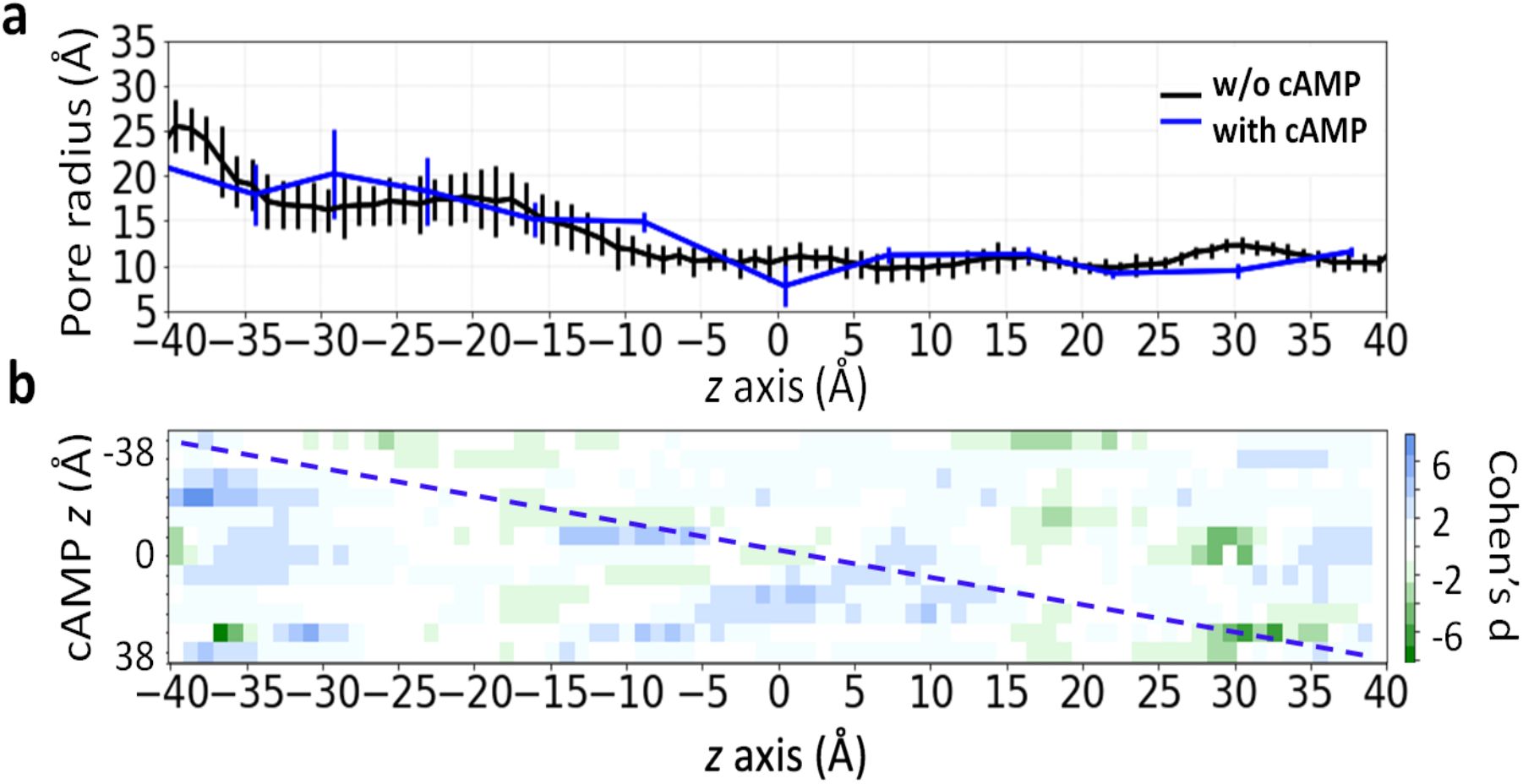
Local and non-local effect of cAMP on Cx26 pore radius. **a**. The mean and standard deviation of Cx26 pore radius as a function of the pore axis (z) during simulations. Black line is the pore radius without cAMP and the blue line is the radius at the position of the cAMP. **b**. Cohen’s d-scores of pore radius distributions (see Eq. 3). The y-axis indicates the z-position of the cAMP, and x-axis is the full-length pore axis. The dashed diagonal line indicates changes in pore radius at the position of the cAMP, while off-diagonal colors indicate cAMP’s non-local effect on the pore radius. Blue color indicates cAMP increase the pore radius, and green color indicates cAMP decrease the pore radius.

What is the cAMP dipole orientation during the voltage induced transition? **Figure 7c** shows scatter plots of dipole vectors of permeating cAMP molecules from the two voltage simulations. Between the intracellular entrance and the major binding site (−40∼-15 Å), dipole angles show orientation under both voltages similar to those at zero voltage, likely due to the large magnitude of the channel local field. While the sampling is scarce in the barrier crossing region (−20<z<20 Å), cAMP shows a clear preference in orientation at z∼0 and at z∼25-30 Å, adopting the opposite orientation under the two voltages. Thus, this position-dependent dipole orientation of the permeant is under the influence of both channel’s internal field and external voltage.

### Position-dependent influence of cAMP on pore radius

The presence of cAMP at different locations inside the pore may have local or non-local effect of the pore radius. To investigate the local effect, we first compared the mean and standard deviations of the pore radius along the length of the pore, without cAMP (**Figure 8a** black), and the radius at the position of the cAMP (**Figure 8a** blue). Notice that these pore radii, obtained using grid-based cavity search program trj_cavity(*25*) ranged between 10 to 15 Å, larger than the 7.5 to 10.5 Å radius obtained using the Hole program (**Figure S1**). This is because the Hole algorithm uses a spherical probe of increasing radius, which underestimates the space within an irregularly shaped pore. It is clear that the influence on pore radius depends where cAMP resides. For instance, the radius increases when cAMP is around z=-10 Å but decreases when it is around z=30 Å, near the extracellular entrance of the pore.

To investigate the non-local effect of cAMP on pore radius, we used Cohen’s d-score of pore radius distributions to compare the mean radius with and without cAMP. **Figure 8b** is a heat map of Cohen’s d-score, which depicts the changes in pore diameter along its length (x-axis of Figure 8b) caused by the presence of cAMP at each z-position (y-axis of Figure 8b). Cohen’s d score was calculated using Eq. 3:

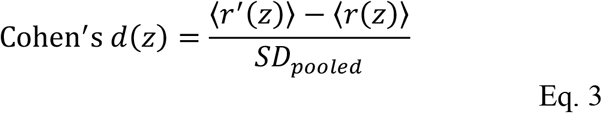

where ⟨*r*(*z*)⟩ is the average pore radius without cAMP (**Figure 8a** black) and ⟨*r*^*^(*z*)⟩ is the average pore radius with cAMP at various locations of the pore, obtained from milestoning simulation trajectories. *SD*_*pooled*_ is the pooled standard deviation of *r*^′^(*z*) and *r*(*z*). For instance, a d score of 6 (blue) means the presence of cAMP increased the mean pore radius by 6*SD*_*pooled*_, and a score of −6 (green) means the presence of cAMP decreased the mean pore radius by 6*SD*_*pooled*_.

On the heatmap, the d score colors along the diagonal line represent changes in pore radius at the position of the cAMP, corresponding to the **Figure 8a** blue profile. The off-diagonal colors represent non-local changes in pore radius away from where the cAMP is in the pore. It is evident that the presence of cAMP can have effects on lumen radius distant from the position of cAMP itself. Most interestingly, the scores on the upper off diagonal are more populated by green color and lower off diagonal shows more blue color. This trend suggests that when cAMP permeates through the pore, it tends to enlarge the radius on the left side (towards intracellular) and narrow the radius on the right side (towards extracellular).

## Discussion

In this work, we use both long-timescale MD simulations and Voronoi-tessellated Markovian milestoning, an enhanced sampling method, to explore how a charged biological signaling molecule, cAMP, permeates a connexin pore. This work builds on our previously developed Cx26 hemichannel model that was validated regarding ionic and molecular permeation properties (*6, 26*). We first obtained the density profile and the inward/outward cAMP flux under +/-200 mV voltages from two 2 μs MD simulations in the presence of multiple cAMPs. These results were compared with the intrinsic potential of mean force (PMF) and the inward/outward mean first passage time (MFPT) of a single cAMP at zero voltage obtained from a total 16.5 μs of multi-replica milestoning. Those two computational approaches – long timescale under voltage (nonequilibrium) and milestoning without voltage (equilibrium) − provided complementary information that allowed detailed analysis of the kinetics of cAMP transit in the absence and presence of voltage. The relation between voltage simulations and milestoning simulations were investigated by deriving the PMF under voltage from the intrinsic PMF, which revealed how mobile ions and protein dipole contribute to the resulting free energy landscape. In addition, unbiased simulations within each milestoning cell allowed us to examine the dipole orientation of cAMP through the pore, with and without voltage, and the short-range and long-range effects of cAMP on pore width.

Both +/-200 mV simulations in 26.5 mM cAMP and single cAMP milestoning simulation featured a prominent intra-pore binding site, characterized by a high density of positive charge in a wide entrance region of the pore. The PMF and MFPT from milestoning further allow us to estimate the binding constants, dissociation and association rates of cAMP inside the channel. Because this site directly communicates with the bulk aqueous compartment, the “saturability” of this site is expected to be more complex than for a single-occupancy ion binding site in a narrow pore. The results also suggest that while more than one cAMP can be within the pore, this multiple occupancy only occurs at this wide binding site, and that there is single cAMP occupancy in the rest of the pore. Thus, we do not anticipate for cAMP the types of multi-occupancy effects seen in many ion channels that involve interactions between sequential single-occupancy binding sites in the permeation path.

At the cAMP concentration used in the voltage simulations, we noted that there were self-interactions among the cAMP molecules at the pore entrance (and not in bulk); cAMP molecules interacted with each other via pi-stacking and hydrogen bonding. This was likely due to the 26.5 mM bulk concentration of cAMP used to obtain a measurable number of transits during the voltage simulation, which was greatly increased at the pore entrance. Although Mg^2+^ ions were included in all simulations, they did not play a role in cAMP clustering or permeation. The cAMP self-interactions are unlikely to occur at any physiological cAMP concentrations. However, this finding does point out a caution when a high concentration of ligands is used in computations to speed up the sampling.

While the cAMP binding sites and barriers are consistent between nonequilibrium simulation and milestoning simulation, the kinetic features are different. Under equilibrium simulation, the cAMP transit time is ∼3 times faster inward than outward. This asymmetric rate is not seen under voltage simulations. By exploring how voltages influence the permeation free energy, we found this is likely due to the negative voltage reducing the free energy barrier for outward cAMP flux. The influence of the voltage of the free energy profile of cAMP permeation was estimated using intrinsic PMF from milestoning and the electrostatic potential change induced by the electric field. The PMF profiles under the two opposite voltages highlight how the external voltage alters the thermodynamics and kinetics of cAMP permeation by changing both the relative free energy as well as free energy barriers. One feature that emerged from this investigation was the recognition of that for a wide pore such as connexin26, the effect of voltage on the mobile charges and polarizable elements within the pore produces changes in the electrostatic field within the pore that affect permeation. These voltage-induced modifications of the reaction field alter the energetic landscape in protein-specific ways. For the connexin channel, these changes are imposed on the intrinsic dipole within the pore, which is responsible for the asymmetric effect of symmetric voltage changes on the overall PMF.

The primary impact of this work will be to establish a way to generate meaningful hypotheses/understanding about the basis of molecular selectivity of connexin channels and to understand how mutations of connexin proteins alter the selectivity and thereby cause human pathologies. Such hypotheses can be tested experimentally and computationally in a synergistic manner. The broader application will be studies of permeation of other biological molecules (e.g., ATP, glutathione, IP3) known to permeate Cx26 channels, and eventually to extend the work to other connexin isoforms as validated atomic models are developed. The methods presented in this study can be applied to understand the molecular permeation through a large pore in general. Input files and raw data used to generate each figure, as well as python3 scripts for milestoning analysis are publicly available at https://github.com/LynaLuo-Lab/Connexin-cAMP-milestoning. Long timescale MD trajectories are publicly available on Anton2 supercomputer.

## Methods

### Force Field and cAMP Parameterization

CHARMM36 force field was used for protein(*27, 28*), POPC lipids(*29*), KCl, and TIP3P water (*30*). Mg^2+^ parameters were from Yoo and Aksimentiev(*31*), in which the van der Waals interaction parameters were fine-tuned to reproduce experimental osmotic pressure. For cAMP, force field parameters were first generated from CHARMM CGenFF(*32*). Additional dihedral fitting between imidazole and pyran groups in the cAMP structure was performed in VMD ffTK plugin(*33*). The final optimized cAMP parameters are provided in **Table S3** and https://github.com/LynaLuo-Lab/Connexin-cAMP-milestoning.

### System Setup and Equilibrium Protocol

The atomistic model of Cx26 was taken from our previous work(*6*). The system of Cx26 embedded in a solvated 1-palmitoyl-2-oleoylphosphatidylcoline (POPC) bilayer with ions and TIP3P water molecules was built and equilibrated following the step-by-step protocol used in Membrane Builder in CHARMM-GUI website(*34, 35*). The terminal amino acid of each segment was capped using acetylated N-terminus (ACE) and methylated C-terminus (CT1), and three disulfide bonds were added between the residue pairs of C53 and C180, C64 and C169, C60 and C174 respectively per protomer, so the entire channel contained a total of 18 disulfide bonds, consistent with the original crystal structure (PDB ID 2ZW3). Two systems containing a single cAMP for milestoning simulations and 27 cAMP for nonequilibrium simulations were constructed (see **Table S1** for system details for Milestoning simulation and Anton2 simulation). The system was energy minimized and serially equilibrated in NVT and NPT ensembles with positional restraints using AMBER18(*36*). Temperature was maintained at 310.15 K using Langevin thermostat (*37, 38*) and 1 atm was maintained by Monte Carlo barostat pressure control (*39, 40*). The time step was 2 fs. Cutoff for calculating van der Waals interactions and short-range electrostatic interactions was set at 12 Å and force-switched at 10 Å. Long-range electrostatic interactions were calculated using the particle mesh Ewald algorithm (*41*).

### Anton2 Simulation Protocol

After 35 ns equilibrium simulation, the system was run on Anton2 supercomputer with 2.0 fs timestep. Lennard-Jones interactions were truncated at 11-13 Å and long-range electrostatics were evaluated using the k-Gaussian Split Ewald method (*42*). Pressure regulation was accomplished via the Martyna-Tobias-Klein (MTK) barostat, to maintain 1 bar of pressure, with a tau parameter of 0.0416667 ps and reference temperature of 310.15 K. The barostat period was set to the default value of 480 ps per timestep. Temperature control was accomplished via the Nosé-Hoover thermostat with the same tau parameter. The mts parameter was set to 4 timesteps for the barostat control and 1 timestep for the temperature control. The thermostat interval was set to the default value of 24 ps per timestep. A 600 ns equilibrium simulation was finished before applying voltage. Constant electric fields of −200 or +200 mV respectively were added for 2μs simulation time with trajectories saved every 200 ps.

### Milestoning MD Simulation Setup

The AMBER18 CUDA version currently does not support Cartesian coordinates as a collective variable. Thus, we pinned two water molecules using high Cartesian restrain with 6000 kcal mol^-1^ Å^-2^ force constant in the top and bottom bulk region (20 Å away from the intracellular entrance and 35 Å away from the extracellular entrance of the channel). We then used the projected distance on the z vector between the nearest pinned water oxygen and center of mass of cAMP to define the Voronoi cells along the z-axis. A cylindrical restraint with a radius of 30Å was applied to cAMP to confine the sampling in the bulk region. To prevent protein drifting, a strong harmonic distance restraint with force constant 2000 kcal mol^-1^ Å^-2^ along *xyz*-axes between the fixed water oxygen in intracellular bulk and the center of mass of the protein was added. The simulation protocol in AMBER18 is the same as above, except all milestoning simulations were run in NVT ensemble. The timestep was 2.0 fs, and each trajectory was saved every 500 frames. Confinement within the Voronoi cells was obtained using flat-harmonic restraint with force constant of 100 kcal mol^-1^ Å^-2^.

### Convergence of PMF and MFPT

A total of 53 Voronoi cells *B*_*i*_ are evenly distributed 2 Å apart along the z-axis, and a milestone state *S*_*ij*_is defined as the boundary between two adjacent Voronoi cells *B*_*i*_ and *B*_*j*_. If *k*_*i*→*j*_is the rate of attempted escape from cells *B*_*i*_ to *B*_*j*_, since at statistical equilibrium the total flux in and out of each cell is zero, thus the equilibrium probability *π*_*i*_ for the system to be in cell *B*_*i*_ satisfies a balance equation:

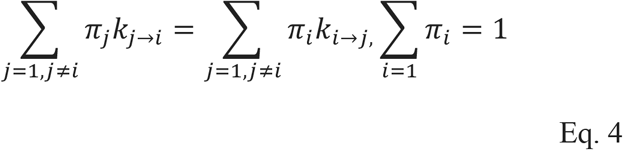

The free energy of each cell can be obtained from the solution of Eq. 4 as -*k*_*B*_*Tl n*(*Π*_*i*_). Running independent simulations in the various cells allow focused sampling on the cells with slow convergence. Here, we monitor the convergence of *π*_*i*_ on the fly by plotting the accumulated rate of attempted escape on both sides of the cell *B*_*i*_, called *k*_*i*→*j*_and *k*_*i*→_*k*, as well as the retention rate inside each cell *B*_*i*_ over time (**Figure 9a**).

**Figure 9.**
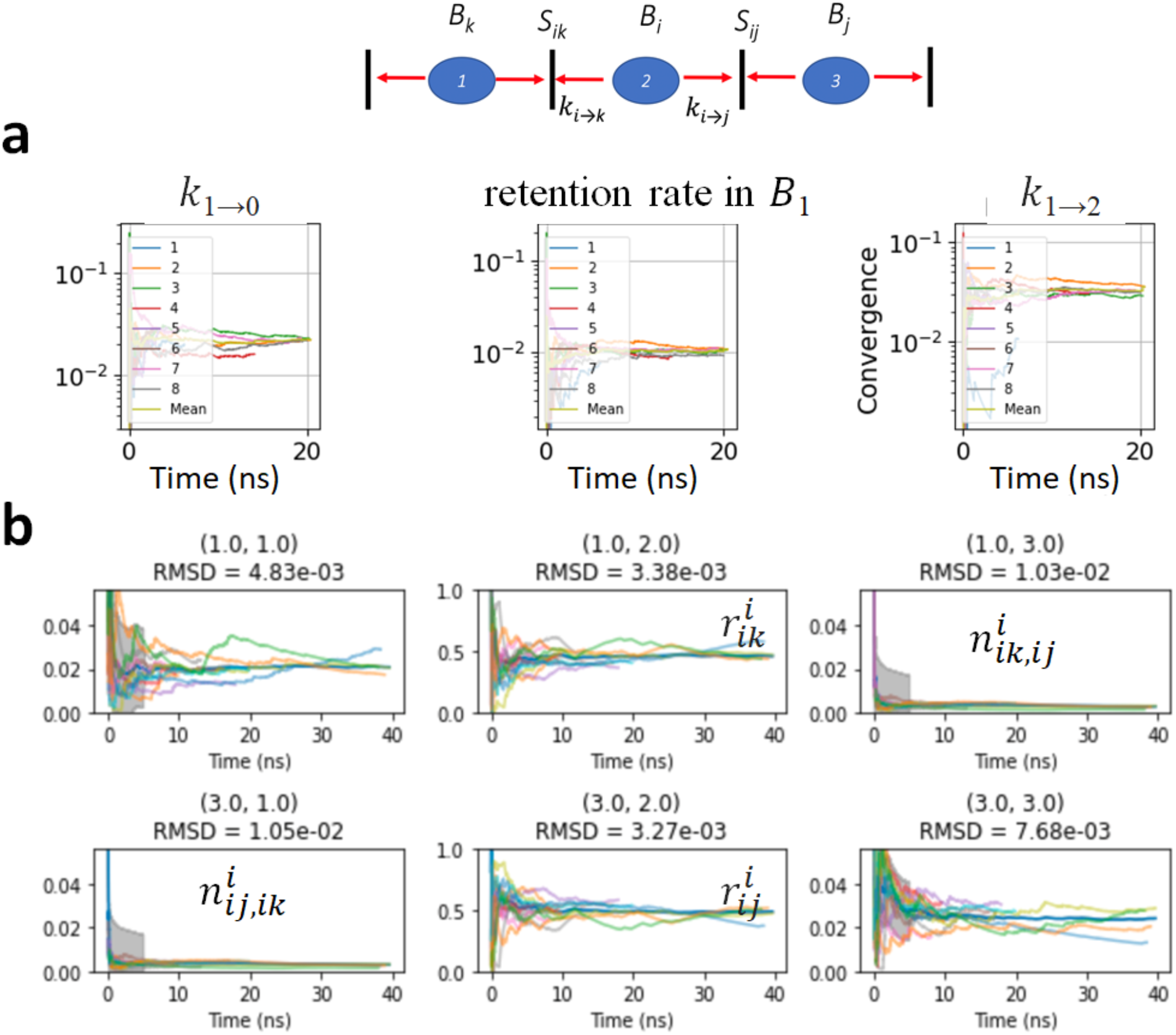
**a**) PMF convergence plots of a Voronoi cell for index 01 (*B*_1_) over time. The left and right plots represent the probability of the attempted escape to the left or right milestone states, *k*_1→0_ and *k*_1→2_. The middle plot is the retention rate inside the cell. Colors represent different replicas and the mean of all the replicas. **b**) convergence plots for MFPT for milestoning cell index 2 (*B*_2_). The upper right and lower left panels correspond to the frequencies of transitions from cell 1 to cell 3, and from cell 3 to cell 1, respectively (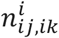 and 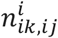). The upper and lower center panels correspond to the percentage of time spent in cell 2 after last touching cell 1 and 3 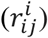, respectively. The other two entries are not used in analysis directly but would correspond to re-entering cell 1 or 3 after last visiting that same cell previously. The final 10 ns (last 10,000 frames) windowed relative RMSD is also given at the top of each panel as a measure of the degree of convergence for the corresponding rate matrix entry components. The convergence plots of all 53 cells are available on Github repository https://github.com/LynaLuo-Lab/Connexin-cAMP-milestoning.

By defining a milestone *S*_*ij*_as the boundary between two adjacent Voronoi cells *B*_*i*_ and *B*_*S*_, the dynamics of the system is reduced to that of a Markov chain in the state space of the milestones indices(*9*). The MFPT between any pair of milestones *S*_*ij*_and *S*_*ik*_ can hence be calculated from the rate matrix whose elements *qij,ik*, the rate of moving from milestone *S*_*ij*_to *S*_*ik*_, are given by:

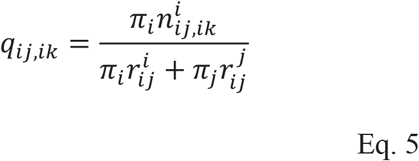

where 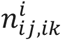 is the number of transitions from *S*_*ij*_to *S*_*i*k_, normalized by the time spend in cell *B*_*i*_, and 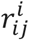 is the time passed in cell *B*_*i*_ after having hit *S*_*ij*_before hitting any other milestone, normalized by the total time spent in cell *B*_*i*_. Therefore, after the convergence of *π*_*i*_, the convergence of MFPT can be monitored directly from the accumulated 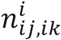 and 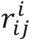 (**Figure 9b**).The final 10 ns windowed relative RMSD is also given at the top of each panel as a measure of the degree of convergence for the corresponding rate matrix entry components. This relative RMSD within 5% (averaged over all replicas) is used as convergence criteria for all Voronoi cells, and in most cases, RMSD is below 2% of the mean value.

## ACKNOWLEDGMENTS

This work was supported by NIH Grant R01-GM130834 and WesternU intramural grant. Computational resources were provided via the Extreme Science and Engineering Discovery Environment (XSEDE) allocation TG-MCB160119, which is supported by NSF grant number ACI-154862. Anton2 computer time was provided by the Pittsburgh Supercomputing Center (PSC) through NIH Grant R01-GM116961. The Anton2 machine at PSC was generously made available by D.E.Shaw Research.

## AUTHOR CONTRIBUTIONS

W.J. prepared the system, re-parameterized cAMP force field, and performed nonequilibrium simulations. Y-C.L. performed milestoning simulations. W.J and Y-C.L analyzed both simulation results. W.M.B-S. prepared milestoning analysis scripts in python3 and performed convergence analysis. W.M.B-S. and L.M. supervised milestoning simulation. Y.L.L, A.H, L.M, and J.C designed the project and wrote the paper with input from all authors.

## Supporting Information

**Figure S1.**
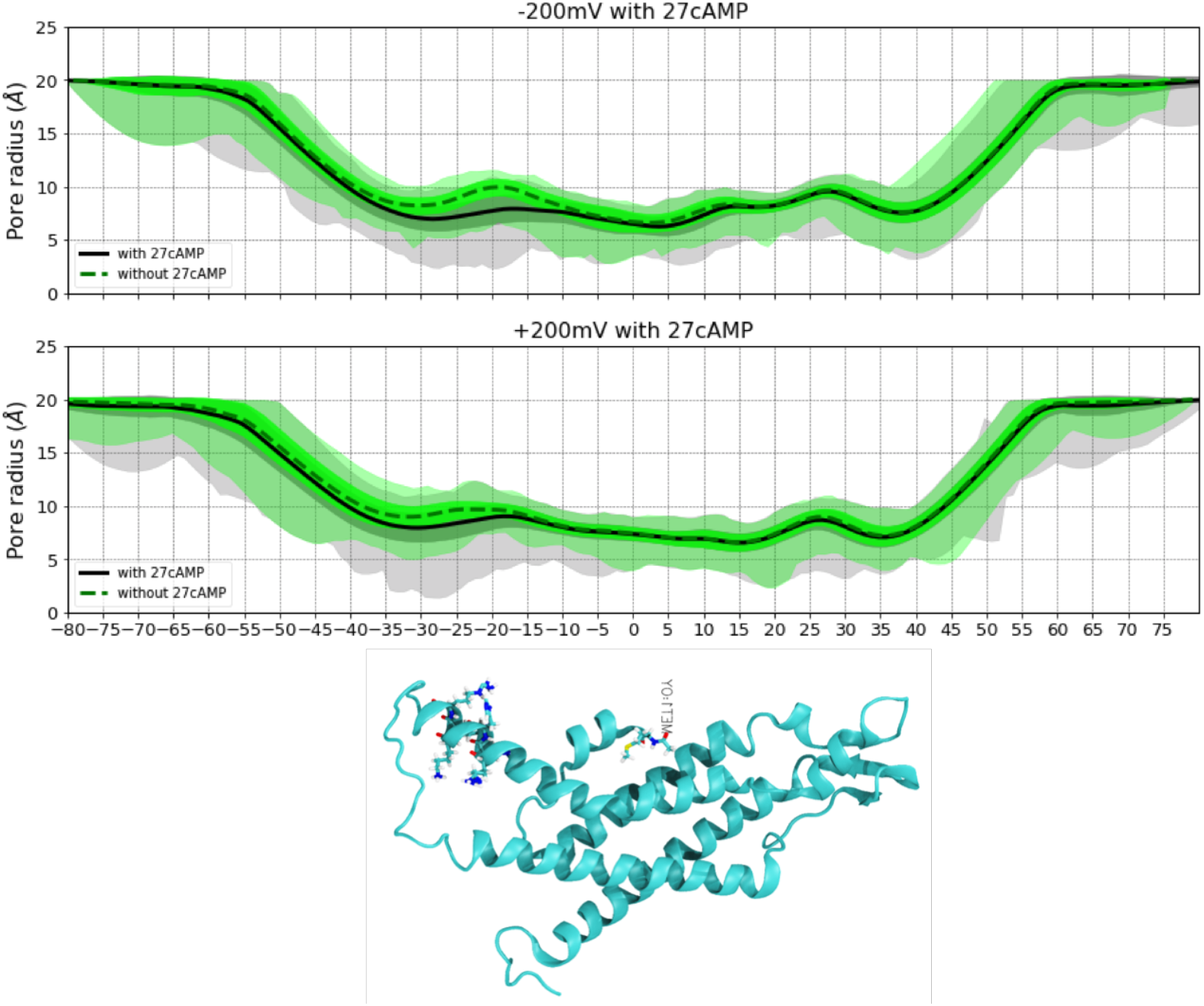
Pore radius profiles from two simulations under +/-200 mV, calculated using Hole program. The black line is the average value over last 1 µs, with 1.2 ns interval between snapshots. The dark grey shade represents the +/-standard deviation, the light grey shade represents the minimum and maximum radius values. The dashed line and green shade are the pore radius profile with protein only.

**Figure S2a.**
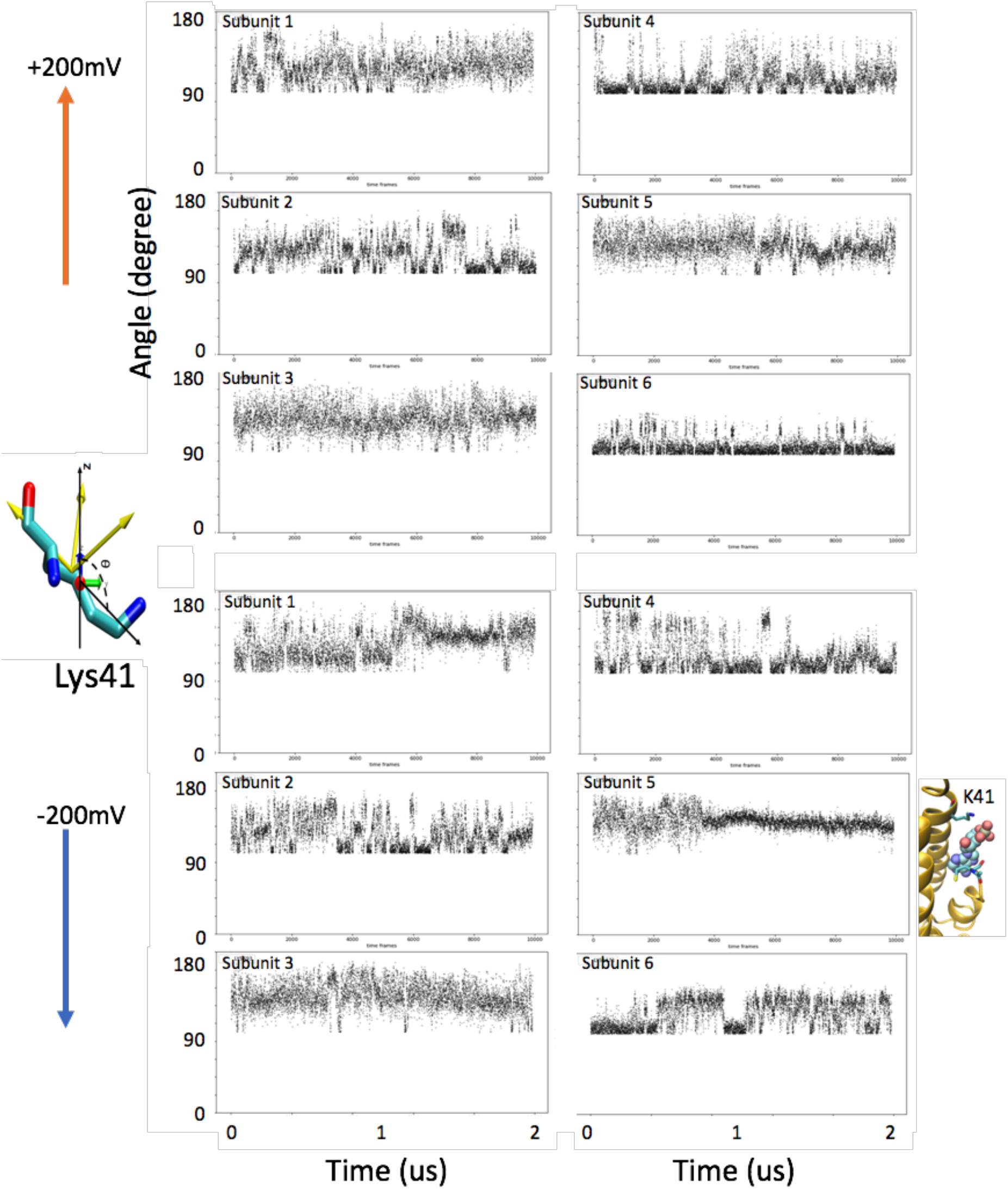
Fluctuation of the angles between the principal axis of Lys41 and z-vector in each subunit under +/-200 mV. Representative Lys residue shown in the licorice model with its principal axes and the angle plotted.

**Figure S2b.**
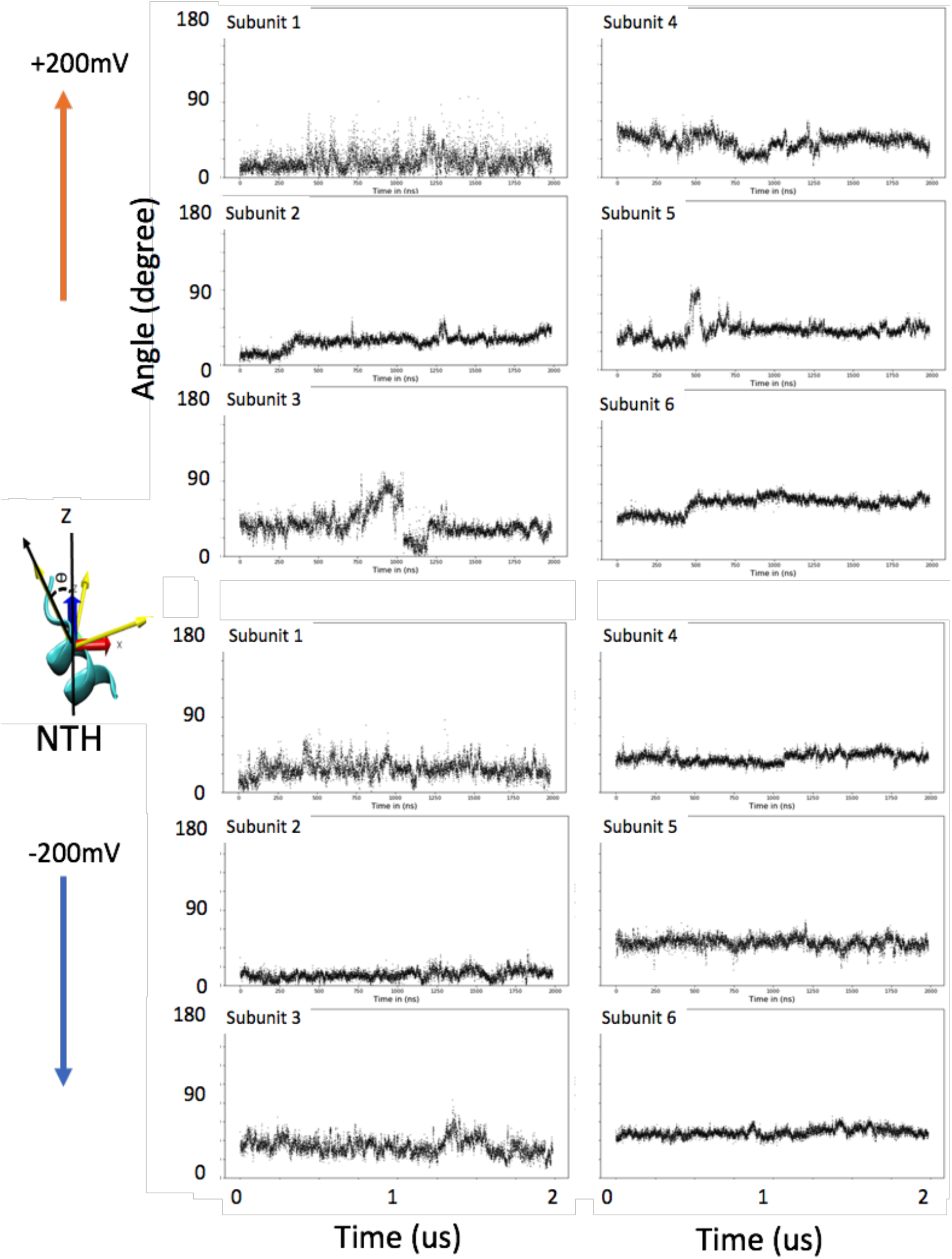
Fluctuation of the angles between the principal axis of NTH (1-11) and z-vector in each subunit under +/-200 mV. Representative NTH shown in the new cartoon model with its principal axes and the angle plotted.

**Table S1.** Description of the computational systems.

|  | Voltage | term patching | Disulfide bonds | Box (Å<sup>3</sup>) | Number (concentration*) |  |  |  |
| --- | --- | --- | --- | --- | --- | --- | --- | --- |
|  |  |  |  |  | atoms | cAMP | K+/Mg2+/Cl- | POPC/water |
| Anton2 | +/- 200mV | ACE CT1 | C53-C180<br>C60-C174<br>C64-C169 | 122×<br>122x<br>154 | 237030 | 27<br>(26.5 mM) | 82/27/163<br>(80/26/160 mM) | 323/56569 |
| Milestoning | 0 mV | ACE CT1 | C53-C180<br>C60-174<br>C64-C169 | 121x<br>121x<br>155 | 234652 | 1<br>(1 mM) | 82/1/137<br>(81/1/135 mM) | 324/56191 |

\*Concentrations were calculated based on number of water molecules.

**Table S2.** Transition time and dwell time (ns) from 2 μs trajectories at +200 mV and −200 mV.

| Transition Event | +200mV (inward flux) |  |  | -200mV (outward flux) |  |  |
| --- | --- | --- | --- | --- | --- | --- |
|  | Transition Time<br>-50<z<50 | Dwell time<br>-50<z<-20 | Barrier crossing<br>-20<z<50 | Transition Time<br>-50<z<50 | Dwell time<br>-50<z<-20 | Barrier crossing<br>-20<z<50 |
| 1 | 222 | 24 | 198 | 160 | 110 | 50 |
| 2 | 1218 | 1058 | 160 | 242 | 210 | 32 |
| 3 | 416 | 308 | 108 | 290 | 50 | 240 |
| 4 | 282 | 248 | 34 | 1496 | 1400 | 96 |
| 5 | 542 | 498 | 44 | 230 | 162 | 68 |
| 6 | 290 | 149 | 140 | 630 | 570 | 60 |
| 7 | 192 | 120 | 72 | 384 | 350 | 34 |
| 8 | 454 | 222 | 232 | 152 | 100 | 52 |
| 9 | 426 | 150 | 276 | 1410 | 1228 | 182 |
| 10 |  |  |  | 198 | 76 | 122 |
| 11 |  |  |  | 390 | 150 | 240 |
| 12 |  |  |  | 445 | 350 | 95 |
| Raw mean | 449 | 308 | 140 | 502 | 396 | 105 |
| Sample mean* | 448 | 305 | 143 | 510 | 398 | 106 |
| 95% CI* | 204-1646 | 149-940 | 79-330 | 249-1571 | 205-1086 | 54-290 |

\*The sample mean and confidence intervals are based on Maximum Likelihood Estimate by fitting the exponential distributions (see code at https://github.com/LynaLuo-Lab/MD-data-uncertainty-analysis/blob/master/confidence_interval_exponential.ipynb).

**Table S3.**
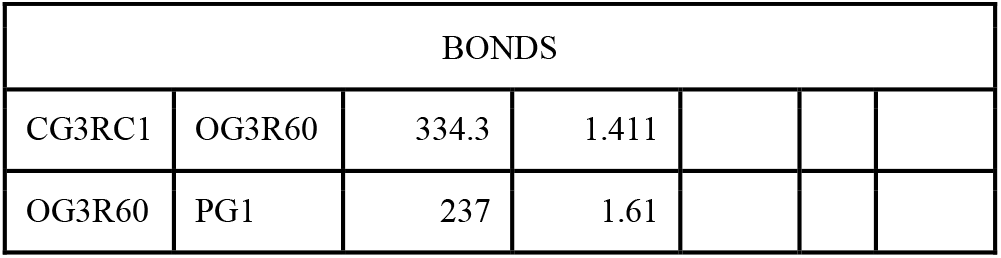
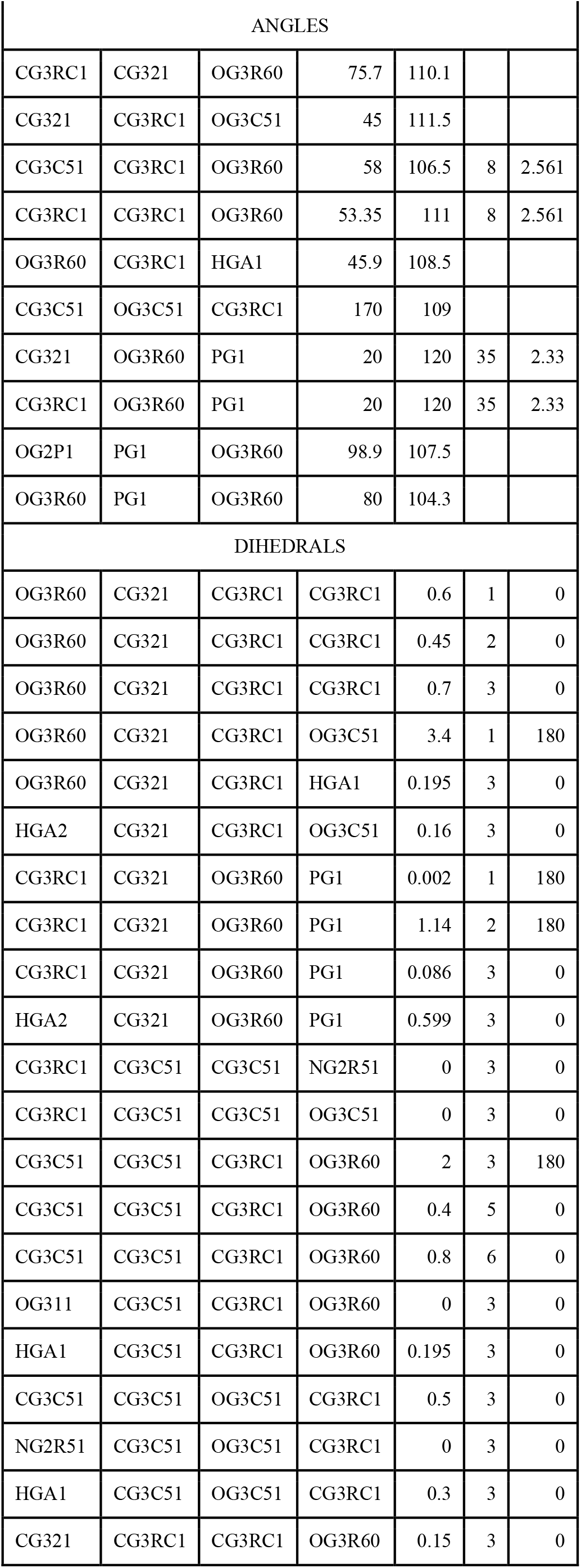
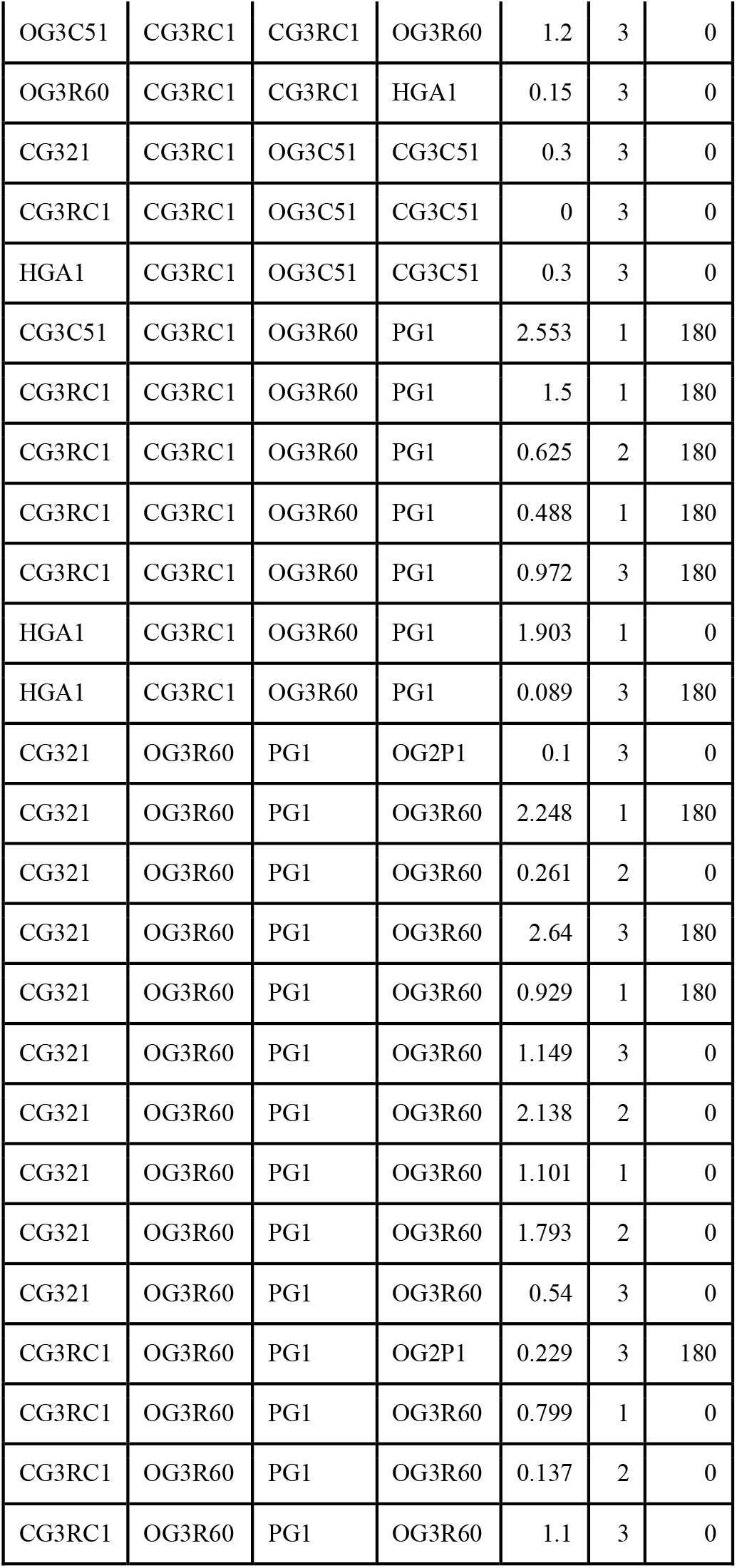
Optimized cgenff force field parameters for cAMP.

